## Supplement for "Eye features and retinal photoreceptors of the nocturnal aardvark (*Orycteropus afer*, Tubulidentata)"

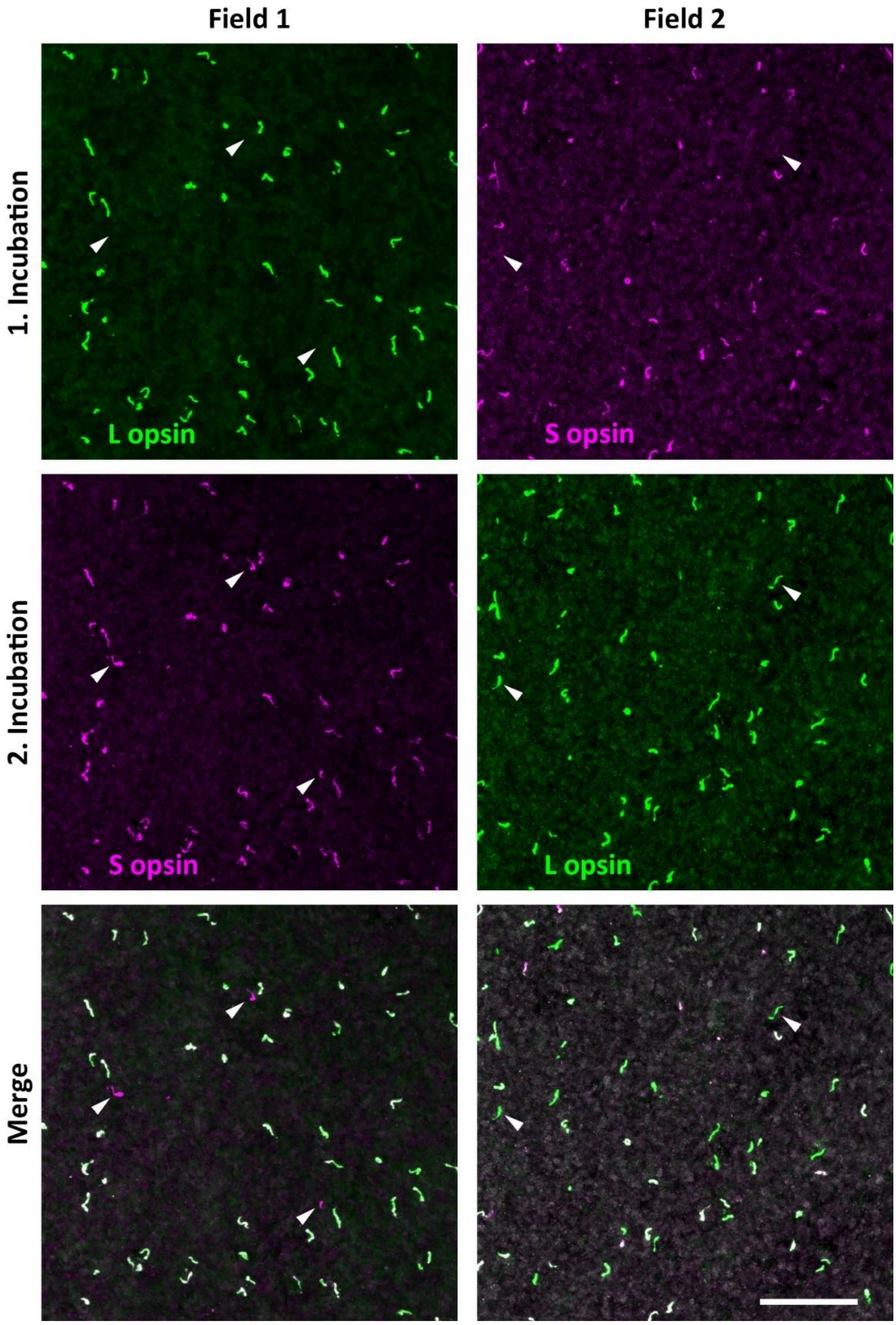


**S1 Fig. Cone photoreceptors**. Sequentially double immunolabelled cones in two neighbouring flat-mounted pieces from an unknown location in midperipheral to peripheral

retina (aardvark 2). The piece of field 1 was first incubated with the rabbit L opsin antiserum JH492, visualized with an Alexa488-coupled secondary antiserum (green). Then it was incubated with the rabbit S opsin antiserum JH455, visualized with a Cy3-coupled secondary antiserum (magenta). The merge shows that all cones are labelled by Cy3, because this secondary antiserum bound to JH455 as well as JH492. The pure S cones are exclusively labelled by Cy3 (arrowheads), all Alexa488-labelled cones contain the L opsin. The neighbouring piece of field 2 was first incubated with the rabbit S opsin antiserum JH455, visualized with the Cy3-coupled secondary antiserum (magenta). Then it was incubated with the rabbit L opsin antiserum JH492, visualized with the Alexa488-coupled secondary antiserum (green). The merge shows that all cones are labelled by Alexa488, because this secondary antiserum bound to JH492 as well as JH455. The pure L cones are exclusively labelled by Alexa488 (arrowheads), all Cy3-labelled cones contain the S opsin. The relative amount of L and S opsin (i.e., labelling intensity) differs between cones. For details of the procedure see Methods. The images are maximum intensity projections of confocal image stacks. Scale bar is 50 µm and applies to all images.

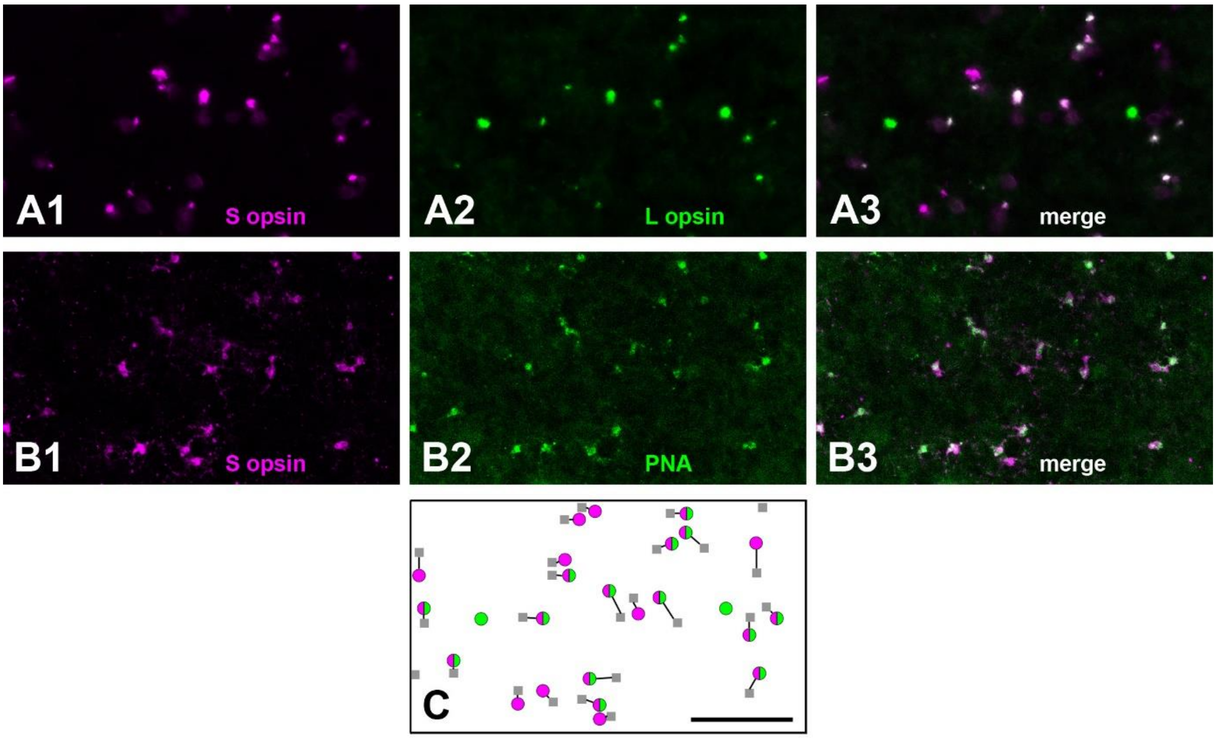


**S2 Fig. Cone photoreceptors and their synaptic pedicles in the outer plexiform layer.** Flat-mounted piece of retina (aardvark 1), triple immunolabelled for S opsin, L opsin and PNA. (**A**) Focus on the outer segments of the opsin-immunolabelled cones; most cones express both opsins (A1, S opsin; A2, L opsin; A3, merge). The field contains two pure L cones. (**B**) Focus on the cone pedicles in the outer plexiform layer; the S opsin-labelled pedicles (B1) are also labelled by PNA (B2), as shown in the merge (B3). (**C**) Schematic illustration of the cones identified in A (S cones, magenta circles; L cones, green circles; dual pigment cones, bipartite circles) and the PNA-labelled pedicles identified in B (grey squares). The pairing between outer segments and pedicles (connecting lines) was checked by following the S opsin-labelled cone axons through the image stacks. The pedicles of the two pure L cones show no PNA label. For two pedicles at the edges of the image, the corresponding outer segments lie outside the frame. Due to faint labelling of some cones, not all cones shown in (C) can be seen in (A) and (B). Scale bar in (C) is 50 µm and applies to all images.
